# Increasing the value of raw bulk milk quality based on mammary glands as production units vs. the udder in dairy cows with mastitis

**DOI:** 10.1101/567271

**Authors:** Gabriel Leitner, Yaniv Lavon, Uzi Merin, Shamay Jacoby, Shlomo E. Blum, Oleg Krifucks, Nissim Silanikove

**Author notes:** **Corresponding Author:** Dr. G. Leitner (G. Leitner).

## Abstract

The current study measured the influence of milk of subclinically infected glands by different bacteria species on the cow’s milk and suggests different parameters for milk payment. The effects of bacterial infection or inflammation on gland milk yield were related to the bacteria species that caused the infection. The volume of milk of the inflamed gland from the cow’s milk yield was significantly lower (*P<0.001*) for the glands previously infected by *Escherichia coli* (PIEc) and those infected with *Streptococcus dysgalactiae.* Coagulation properties, rennet clotting time (RCT) and curd firmness (CF) also depended on the bacteria causing the infection. RCT values of all the inflamed glands were significantly longer (*P<0.001*) and CF values were significantly lower than that of the healthy ones. Moreover, in the whole milk, CF was also significantly lower and not proportional to the volume of the milk from the inflamed gland of the cow’s milk. Calculating the predicted 40% dry matter curd weight (PCW) on the cow level, including the healthy and inflamed glands or the healthy glands alone, found that for 9 of 13 PIEc cows, the presence of the affected gland’s milk in the whole cow milk resulted in a negative PCW value. Likewise, 5 of 20 cows infected by *S. dysgalactiae* had negative delta values. Unlike the latter bacteria, PCW from milk of glands infected with CNS increased, although in a lower magnitude than in the healthy glands. No correlation was found between logSCC in the whole cow milk (healthy and inflamed glands) and PCW.

## Introduction

Mastitis in dairy cows remains one of the major issues of animal welfare in dairy farms [1]. It is also a major cause of economic losses in dairy production, due to culling of cows and increased replacement costs, treatment costs and discarding of abnormal milk [2,3]. Addition of milk from inflamed (mastitic) mammary glands, with or without detected bacteria, to the bulk milk tank, a routine practice in most cases of subclinically affected cows, reduces the whole tank’s milk quality [4,5]. Usually, emphasis is given to clinical mastitis, because it is notable and demands immediate response by the farmer. During clinical infection, milk is usually altered significantly and therefore it is discarded, and if antibiotic treatment is practiced, the milk is discarded until the disappearance of antibiotic residues. Thus, the additional cost of treatment and the economic loss per clinical event is ∼$300-400 [6-8]. Genetic selection for infection resistance and preventing mammary infections through improved management with emphasis on the milking parlors result in very low improvement due to increased milk production and the intensiveness of the dairy industry.

The questions regarding subclinical and post clinical infection are complicated because: 1. Diagnosis requires specific tests such as somatic cells count (SCC) or California mastitis test (CMT) accompanied by bacteria isolation; 2. Routinely, milk recording test and/or on-line information or data are collected on the cow level and not on every gland separately. For the farmer, the cow is the milk producing unit, while for the dairy that buys the milk, it is the bulk milk tank. Regularly, milk recording test provides information on the milk produced and secreted from the four glands, thus, the milk of the inflamed gland is diluted by the milk from the other healthy glands. If individual/bulk milk SCC is the criteria for milk suitable for the industry, then the low-quality milk from the inflamed glands is ignored. Thus, many farmers ignore cows with SCC of 5×10^5^ cells/mL and 1-2×10^6^ cells/mL if no clinical symptoms and/or changes in color and curd flakes are visible in the milk. Moreover, in cases of clinical mastitis, mainly by Gram negative bacteria, the cows return to normal milking after the antibiotic residues are cleared and no visible signs of flakes in the milk occur. Previous studies showed that milk from infected glands disturbs curd formation and decrease cheese yield and these decreases are related to the bacteria causing the infection and not to the number of somatic cells [9-11]. Moreover, in cases of *Escherichia coli* infection, the milk’s low quality can persist for weeks after the eradication of the bacteria [12]. Depending on country, the dairy industry pays for milk volume according to certain formula, which include fat and protein levels, bacteria count and SCC [13]. In many countries, bulk milk tank SCC (BMTSCC) < 2×10^5^ cells/mL is considered high quality milk that can be payed bonuses. However, BMSCC is the average of milk from all the individual cows glands that were milked into the bulk tank, therefore, in each bulk tank the same SCC can be the result of milk with different qualities and SCC levels of the individual cows, which is the combination of all the glands. The various effects of SCC levels in raw milk on cheese yield and quality are tabulated in [11] and in a recent study, that did not show differences in cheese yield made from bulk milk if its SCC was 2×10^5^ or 3×10^5^ cells/mL [5].

The aim of the current study was: To measure the influence of the milk of subclinically infected glands by coagulase negative staphylococci (CNS) or *Streptococcus dysagalactiae* or post infection by *E. coli* on the cow’s milk and suggest different parameters for milk payment.

## Materials and Methods

### Animals

Animal experiments were approved by the ARO Committee of Animal Experimentation and followed its ethical guidelines. Forty Israeli Holstein cows at the Agricultural Research Organization, the Volcani Center, Bet Dagan were enrolled to the study. The average milk yield in the herd was >11,500 L during 305 days of lactation. Animals were fed a typical Israeli total mixed ration containing 65% concentrate and 35% forage (17% protein). Food was offered *ad lib* in mangers located in sheds. Cows were milked thrice daily (05:00, 13:00 and 20:00) in a dairy parlor equipped with an on-line computerized AfiFarm Herd Management data acquisition system that includes the AfiLab milk analyzer (Afimilk, Afikim, Israel). All cows were identified with one chronically (40-140 d) infected mammary gland with CNS (n=7) and *S. dysagalactiae* (n=20) or post clinical infection with *E. coli* (PIEc), which were 17-40 d after detection of the bacterium (n=13). The infected and the uninfected (healthy) glands (with no bacteria finding) were tested and confirmed by 3 consecutive sampling, including bacteriology and milk composition, including SCC.

### Sample collection and analyses

When a cow was identified as suited for the study (infected in a single gland with identified bacteria and elevated SCC or inflamed - PIEc) it was enrolled to the study. At noon milking, the infected gland was milked into a separate container and the other 3 healthy glands into another, and milk volume was recorded. From the two containers 0.5-1.0 L was sampled for future analyses. Then, the milk of the infected/inflamed gland was mixed with the rest of the milk (whole cow milk) and a third milk sample was taken. The three samples were analyzed for: SCC with the Fossomatic 360 (Foss Electric, Hillerød, Denmark) and gross milk composition, i.e., protein, casein, fat and lactose contents with the Milkoscan FT6000 (Foss Electric). Analyses were performed at the Israel Cattle Breeders Association Laboratory (Caesarea, Israel). Milk coagulation time (RCT) and curd firmness (CF) were tested by the Optigraph^©^ (Ysebaert, Frepillon, France) using Fromase 15 TL (Gist-Brocades nv, Delft, The Netherlands) diluted (1:100) to achieve clotting within ∼900 s, as described in detail by Leitner et al. [10]. Leukocytes differentiation was performed by flow cytometry (FACs Calibur, Becton-Dickinson, San Jose, CA, USA) as described by Leitner et al. [14]. Tests were preforming within 24 h after samples collection with milk stored at 4 °C.

### Calculation of the expected curd firmness of the whole cow’s milk and the correlation between curd firmness and 40% dry matter cheese weight at 24 h

In order to calculate the contribution (plus or minus) of the milk of the inflamed gland to curd produced from the milk, the following set of equations was used:

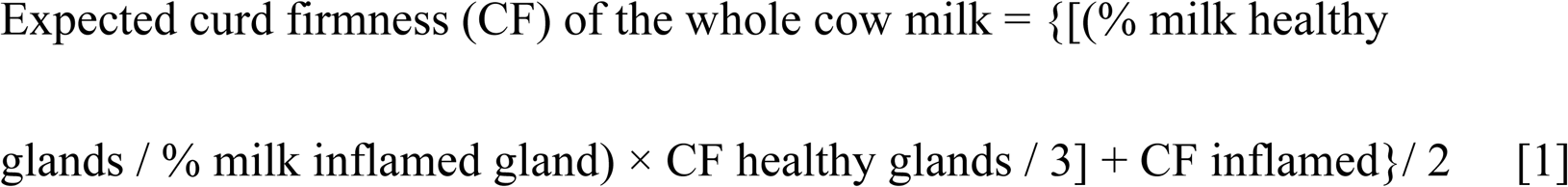

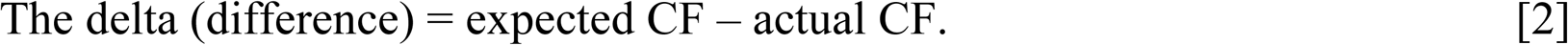

Cheese making at the laboratory was performed as described in detail by Katz et al. [15]. From the data obtained, a correlation equation of curd weight at 24 h and CF was fitted (Supplementary Figure).

From the regression line a set of equations were written in order to calculate the expected curd yield of the cow’s milk.

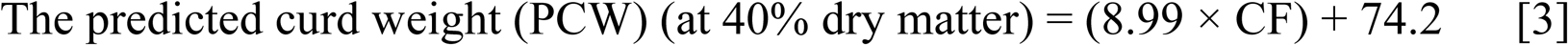

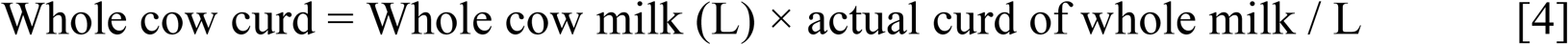

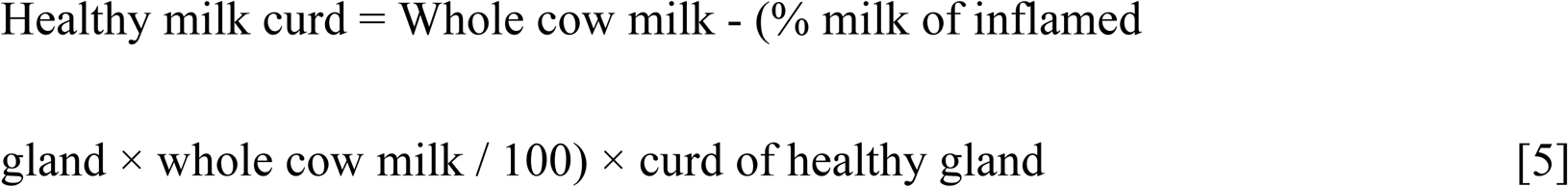

### Statistical analyses

This experiment tests the differences between the dependent variables at the gland level of different cows infected by different bacteria. The analyses were carried out using the GLM procedure of SAS [16] with the general form: Dependent variable = bac + gland + bac × gland + error. Where: bac = three different bacteria causing mastitis: *E. coli, Streptococcus*, Coagulase negative staphylococci, gland = three levels within a cow: healthy, infected and mix (sample that were taken from infected and healthy glands and mix together) and interactions between bacteria and different gland. The dependent variables were: LSCC, fat (% of healthy), fat (g/L), protein (% of healthy), protein (g/L), lactose (% of healthy), lactose (g/L), CD18 (%), PMN (%), CD4 (%), CD8 (%), M Ø(CD14), RCT (sec), RCT (% of healthy), CF (V), CF (% of healthy).

To compare levels within a variable, we ran the Bonferroni adjustment for multiple comparisons. Data are presented as means and SEM.

## Results

Cows were in different lactations (1-6), days in milk (14-463) and milk yield ranged from 21-60 L/day. The values related to the latter three parameters according to bacteria type were: PIEc 3.6±0.2, 175±24, 41.8±2.0; *S. dysgalactiae* 2.4±0.1, 151±14, 46.6±1; CNS 2.7±0.3, 134±13, 47.6±1.8, respectively. The effects of bacterial infection or the inflammation on gland milk yield were related to the bacteria that caused the infection. In calculating the milk volume of the inflamed gland from the cow’s milk yield on the test day, the assumption was that each of the 4 glands contribute 25%, regardless of its position - front or rear. Percent daily milk yield of individual inflamed glands from the total cow’s yield according to the infecting bacteria, including the mean of all the gland (horizontal dotted lines), are presented in Fig. 1. Milk production of the PIEc glands or of those infected with *S. dysgalactiae* was significantly lower (*P<0.001*) than the healthy ones and significantly different between the two species (16.2% vs. 21.1%, respectively). Milk yield of the glands infected with CNS was lower but did not differ significantly from the healthy glands (Fig. 1).

**Figure 1.**
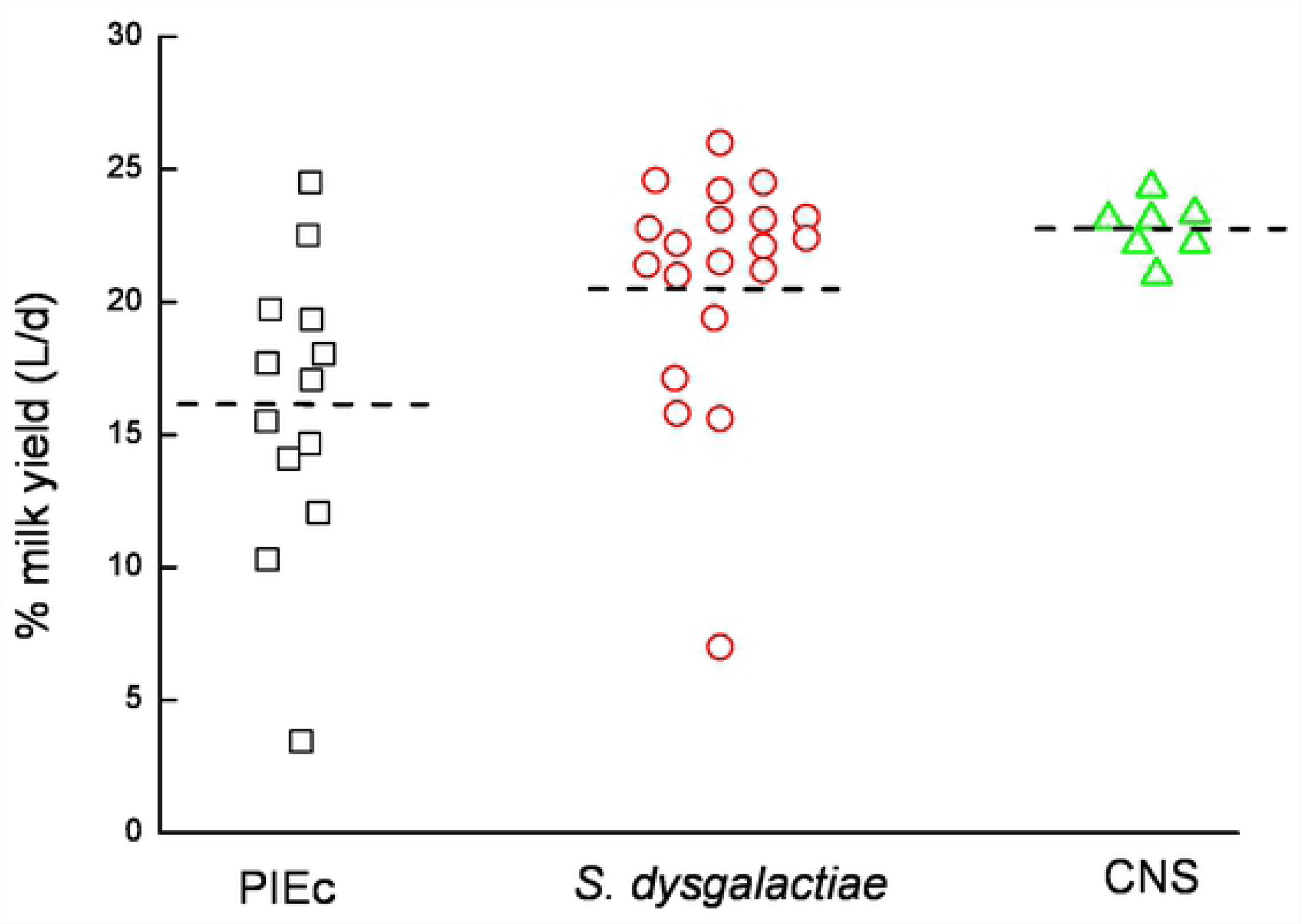
Percent daily milk yield of individual infected glands from the total cow’s yield, according to the infecting bacteria, including the mean of all the gland (horizontal dotted lines) (PIEc – post infected by *E. coli*; *S. dysgalactiae – Streptococcus dysgalactiae*; CNS – coagulase negative staphylococci).

The effects of bacterial infection or the inflammation on gross milk composition, SCC and its differentiation and coagulation properties were significantly different and were related to the bacteria causing of the infection. Milk of the PIEc glands contained significantly lower levels (*P<0.001*) fat and lactose and higher protein than the milk of healthy glands (Table 1). Milk of *S. dysgalactiae* infected glands contained significantly lower (*P<0.001*) lactose and higher protein than milk of healthy glands (Table 1). Milk of the CNS infected glands contained significantly lower level (*P<0.001*) of lactose and higher fat than milk of healthy glands. In the whole cow’s milk (the mixture of healthy glands and the inflamed one), the levels of gross milk composition were proportional to the volume of milk from the infected glands out of the cow’s milk. Changes in coagulation properties, RCT and CF, also depended on the bacteria causing the infection. RCT values of all the inflamed glands were significantly longer (*P<0.001*) than that of the healthy ones (Table 2). Moreover, RCT of the whole milk was also significantly longer and not proportional to the volume of the milk from the inflamed gland of the cow’s milk. The CF values of all the inflamed glands were significantly lower than those of the healthy ones (Table 2). Only 4 of 13 milk samples from the PIEc cows and 7 of 20 of the *S. dysgalactiae* infected ones coagulated (Figs. 2A, B). All the milk of the glands infected with CNS coagulated, but with lower CF values (Fig. 2C). In the whole milk, CF was also significantly lower and not proportional to the volume of the milk from the inflamed gland of the cow’s milk (Table 2). Among the three bacteria, the CF values of the whole milk of PIEc and *S. dysgalactiae* were significantly lower (*P<0.001*) than the calculated value according to the milk volume of the infected glands from the cow’s milk yield and the CF values of the inflamed and healthy ones (Eq. 1). The mean CF of the whole milk from the calculation of PIEc was −57.8% and from *S. dysgalactiae* −11.5% (Fig. 3; Eq. 2). Moreover, of the whole cow’s milk, 3 samples from PIEc and 3 infected with *S. dysgalactiae* did not coagulate (Figs. 2A, B).

**Table 1.** Gross milk composition of uninfected (Healthy), infected and the whole cow’s milk (mixture of healthy glands and the infected one) and the percent change from the healthy glands (Reference) according to infecting bacteria. Values within rows with no common superscript differ significantly (*P* < 0.05).

|  |  | Fat |  | protein |  | Lactose |  |
| --- | --- | --- | --- | --- | --- | --- | --- |
|  |  | g/L | % of Healthy | g/L | % of Healthy | g/L | % of Healthy |
| PIEc | Healthy | 32.0±3.4<sup>a</sup> | Reference | 35.0±1.7<sup>c</sup> | Reference | 52.2±0.5<sup>a</sup> | Reference |
|  | Inflamed | 23.6±4.7<sup>b</sup> | -23.4±11.8<sup>a</sup> | 39.3±2.9<sup>a</sup> | +12.4±5.6<sup>a</sup> | 40.3±2.6<sup>c</sup> | -21.1±4.9<sup>a</sup> |
|  | Whole | 30.3±3.5<sup>a</sup> | -6.2±2.8 | 36.2±1.9<sup>bc</sup> | +3.5±1.7 | 47.9±1.5<sup>b</sup> | -6.4±2.5 |
| Strep. | Healthy | 25.8±2.2 | Reference | 31.9±0.8<sup>c</sup> | Reference | 52.3±0.4<sup>a</sup> | Reference |
|  | Infected | 25.5±1.9 | -1.8±6.3 | 35.6±0.9<sup>a</sup> | +11.8±2.2<sup>a</sup> | 43.5±1.5<sup>b</sup> | -16.6±3.0<sup>a</sup> |
|  | Whole | 25.7±2.0 | -0.7±4.2 | 32.6±0.8<sup>bc</sup> | +2.4±1.3 | 50.3±0.7<sup>a</sup> | -13.8±5.0 |
| CNS | Healthy | 29.3±2.5 | Reference | 32.5±1.0 | Reference | 54.4±0.5<sup>a</sup> | Reference |
|  | Infected | 33.7±6.0 | +20.1±28.8 | 32.8±1.0 | +1.2±3.2 | 46.9±2.8<sup>b</sup> | -3.8±5.0<sup>a</sup> |
|  | Whole | 29.4±2.4 | +0.4±2.1 | 32.7±1.0 | +0.7±0.6 | 53.7±0.7<sup>a</sup> | -1.3±0.7 |

PIEc – post infected by *E. coli*
Strep. – *S. dysgalactiae*
CNS – coagulase negative staphylococci

**Table 2.**
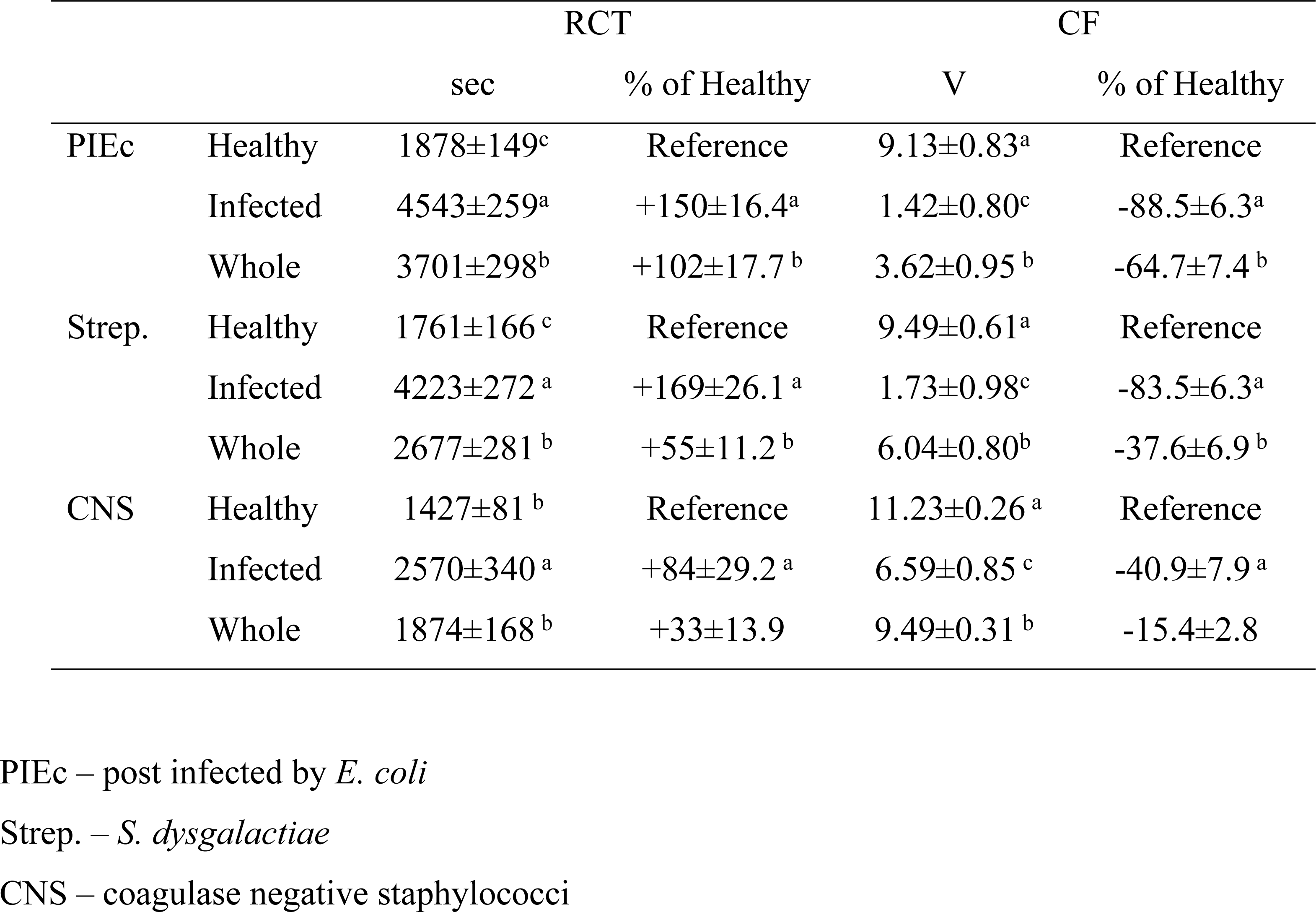
Coagulation properties (curd firmness - CF and rennet clotting time - RCT) of uninfected (Healthy), infected and the whole cow’s milk (mixture of healthy glands and the infected one) and the percent change from the healthy glands (Reference) according to infecting bacteria. Values within rows with no common superscript differ significantly (*P* < 0.05).

|  |  | RCT |  | CF |  |
| --- | --- | --- | --- | --- | --- |
|  |  | sec | % of Healthy | V | % of Healthy |
| PIEc | Healthy | 1878±149<sup>c</sup> | Reference | 9.13±0.83<sup>a</sup> | Reference |
|  | Infected | 4543±259<sup>a</sup> | +150±16.4<sup>a</sup> | 1.42±0.80<sup>c</sup> | -88.5±6.3<sup>a</sup> |
|  | Whole | 3701±298<sup>b</sup> | +102±17.7<sup>b</sup> | 3.62±0.95<sup>b</sup> | -64.7±7.4<sup>b</sup> |
| Strep. | Healthy | 1761±166<sup>c</sup> | Reference | 9.49±0.61<sup>a</sup> | Reference |
|  | Infected | 4223±272<sup>a</sup> | +169±26.1<sup>a</sup> | 1.73±0.98<sup>c</sup> | -83.5±6.3<sup>a</sup> |
|  | Whole | 2677±281<sup>b</sup> | +55±11.2<sup>b</sup> | 6.04±0.80<sup>b</sup> | -37.6±6.9<sup>b</sup> |
| CNS | Healthy | 1427±81<sup>b</sup> | Reference | 11.23±0.26<sup>a</sup> | Reference |
|  | Infected | 2570±340<sup>a</sup> | +84±29.2<sup>a</sup> | 6.59±0.85<sup>c</sup> | -40.9±7.9<sup>a</sup> |
|  | Whole | 1874±168<sup>b</sup> | +33±13.9 | 9.49±0.31<sup>b</sup> | -15.4±2.8 |

PIEc – post infected by *E. coli*
Strep. – *S. dysgalactiae*
CNS – coagulase negative staphylococci

**Figure 2a.**
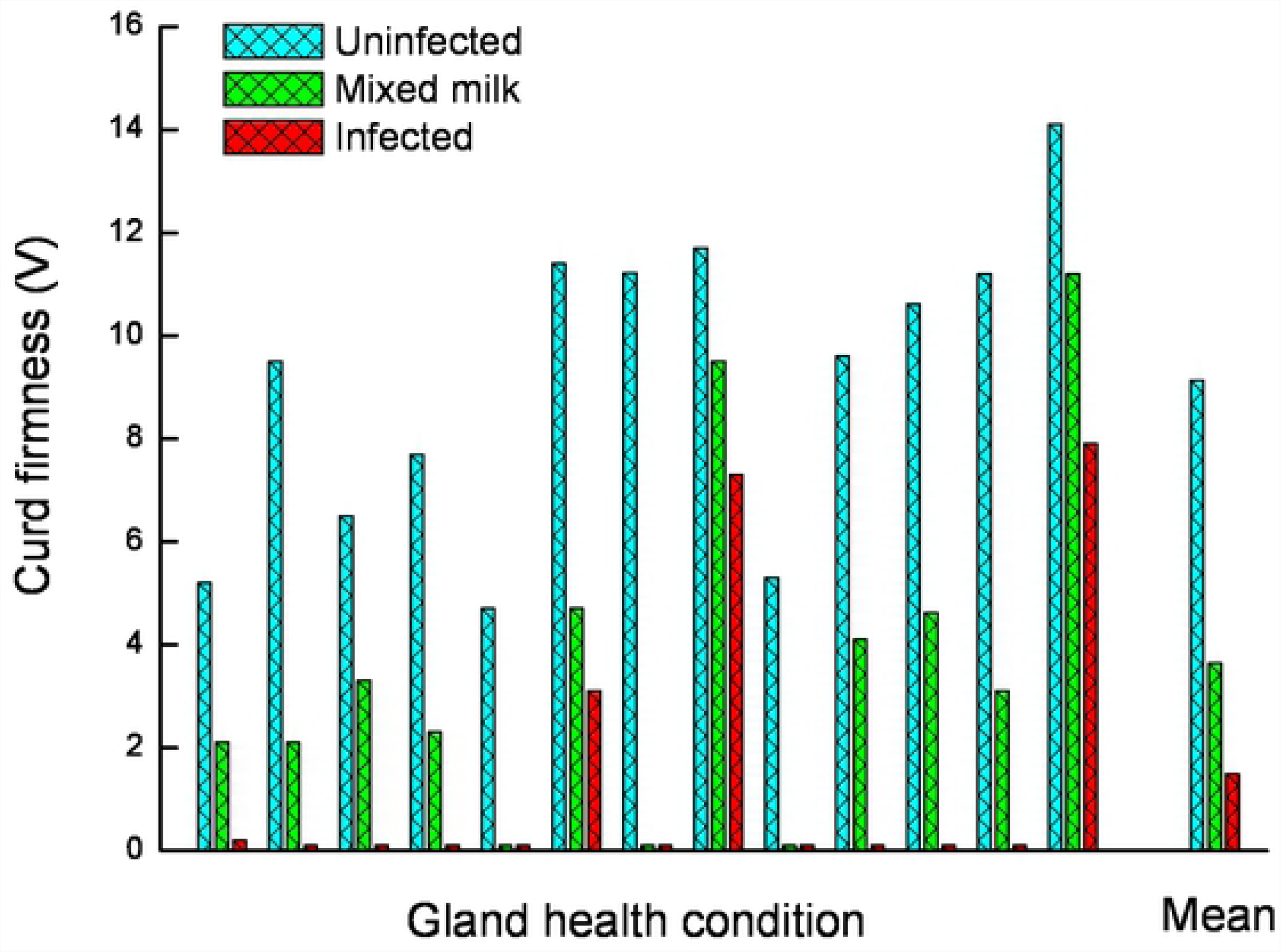
Curd firmness (CF) of individual milk samples of healthy, whole milk and glands post infected by *E. coli* (PIEc), including the mean CF value.

**Figure 2b.**
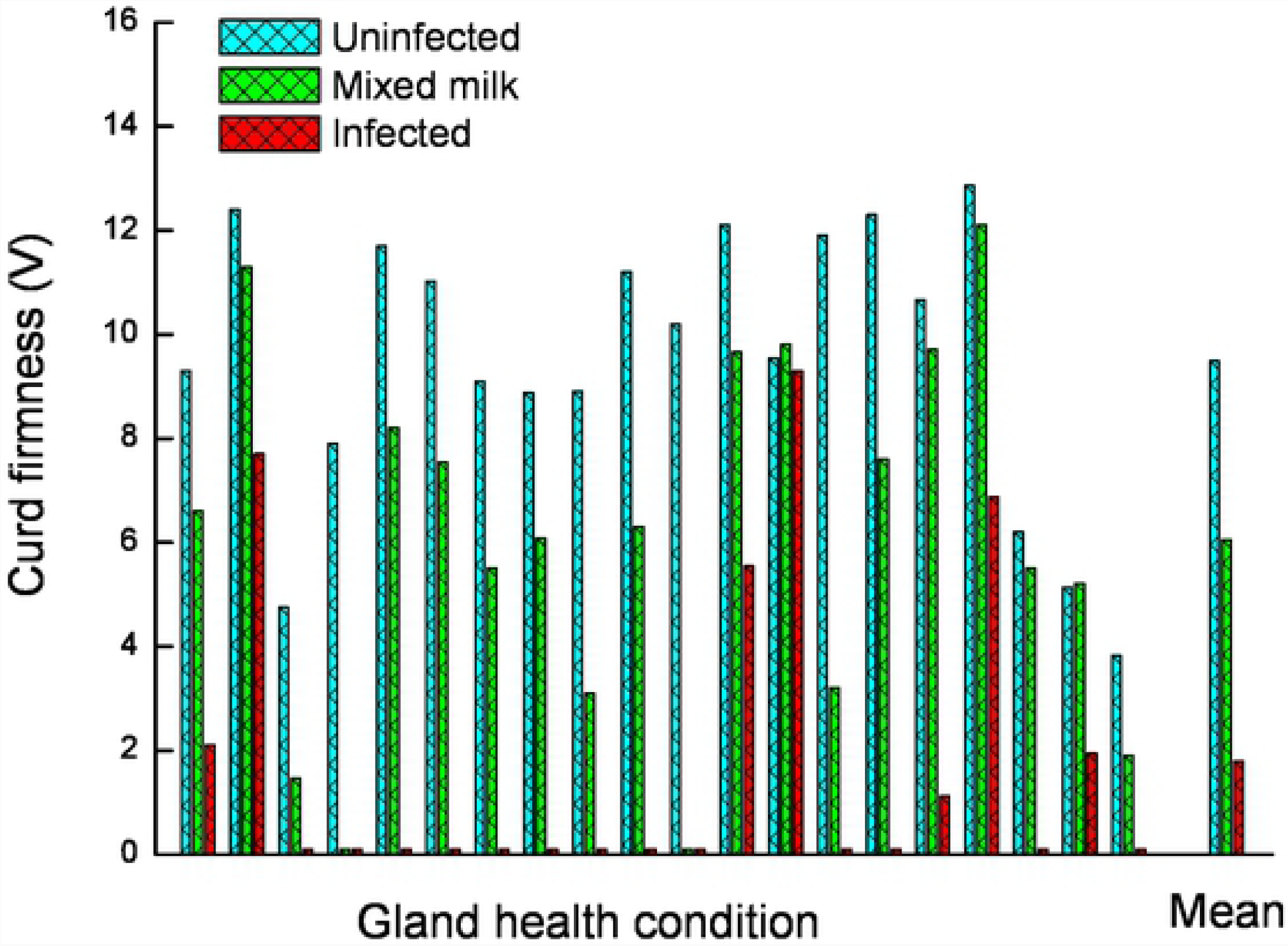
Curd firmness (CF) of individual milk samples of healthy, whole and glands infected by *Streptococcus dysgalactiae*, including the mean CF value.

**Figure 2c.**
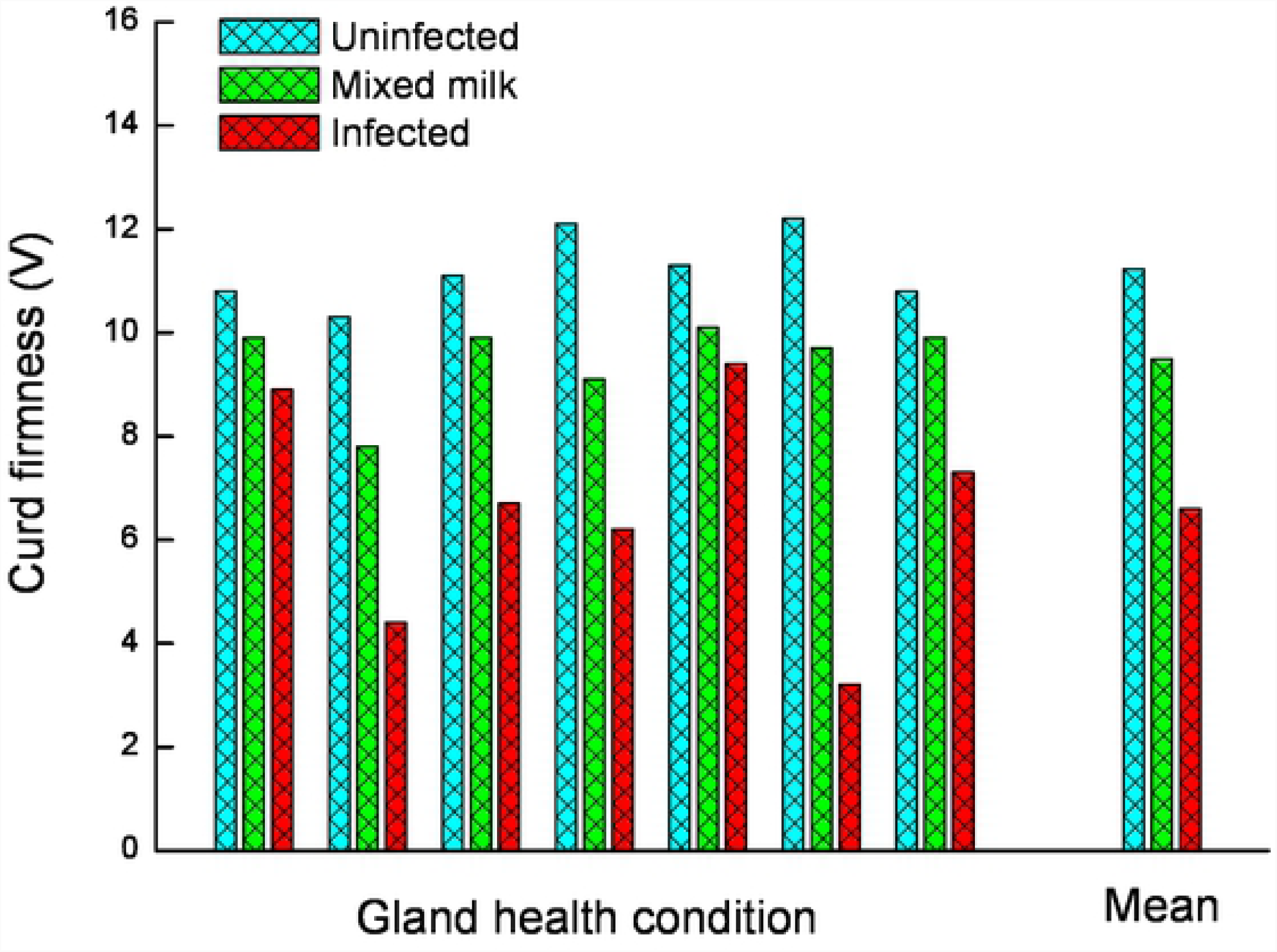
Curd firmness (CF) of individual milk samples of healthy, mixed and glands infected by coagulase negative staphylococci, including the mean curd firmness value.

**Figure 3.**
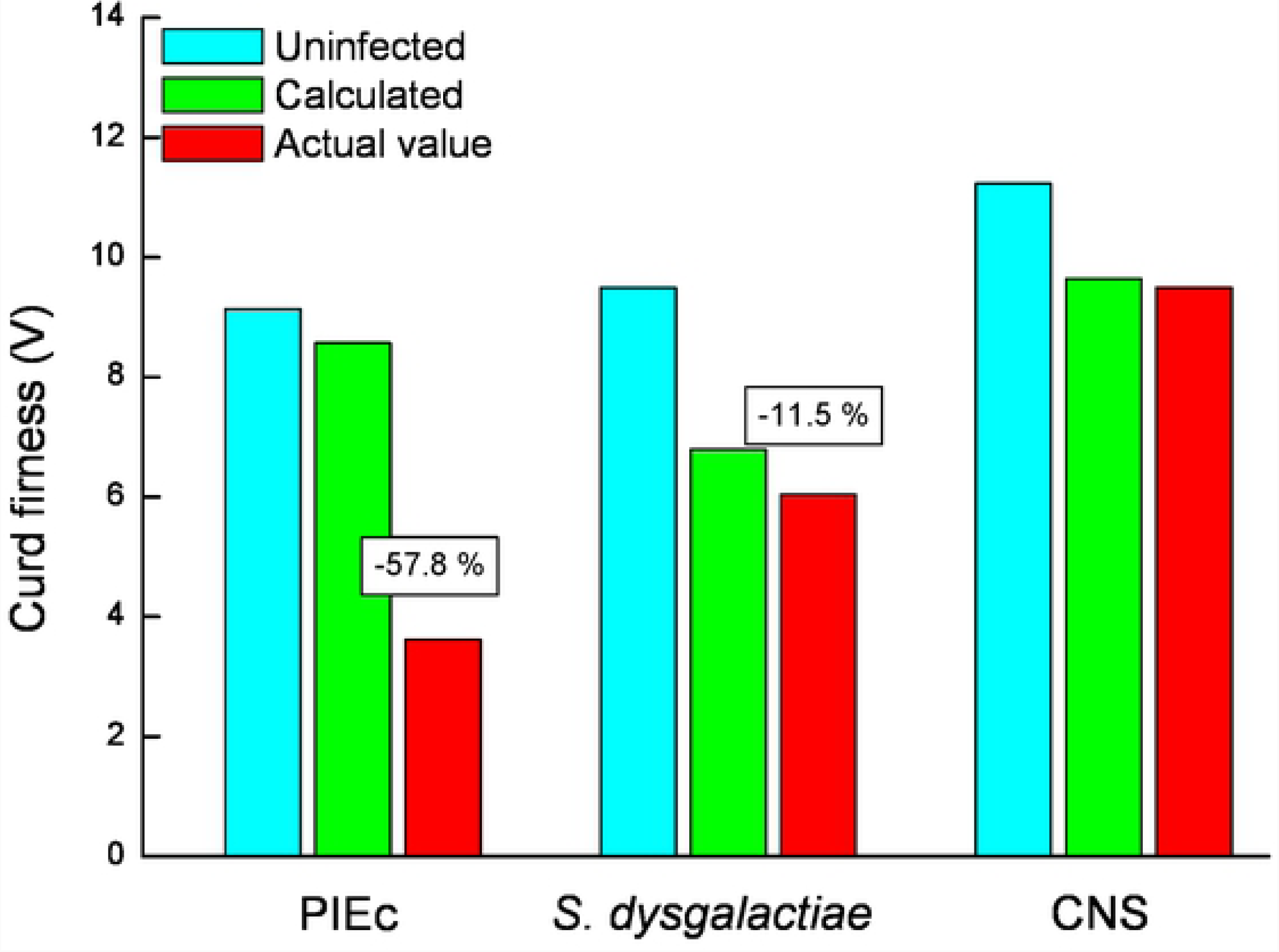
Curd firmness (CF) of milk of healthy glands and the calculated and actual CF value according to the mixed milk volume (Eq. 5). Numbers over bars indicate the difference between the calculated and the actual CF value (PIEc – post infected by *E. coli*; *S. dysgalactiae – Streptococcus dysgalactiae*; CNS – coagulase negative staphylococci).

The predicted 40% dry matter curd weight (PCW) (Eq. 3) on the cow level, including the milk of the healthy and inflamed glands (Eq. 4), the milk from the healthy glands alone (Eq. 5) and the difference between the two for each cow and bacteria are summarized in Fig. 4. In 9 of 13 PIEc cows, the presence of the affected gland’s milk in the whole cow milk resulted in a negative PCW value, higher than the expected if the milk from the inflamed gland had been discarded. Likewise, 5 of 20 cows infected by *S. dysgalactiae* had negative delta values. Unlike these bacteria, the milk from the glands infected with CNS, the PCW increased, although in a lower magnitude than in the healthy glands. No correlation was found (R^2^= 0.4) between logSCC in the whole cow milk (healthy and inflamed glands) and PCW (Fig. 5).

**Figure 4a.**
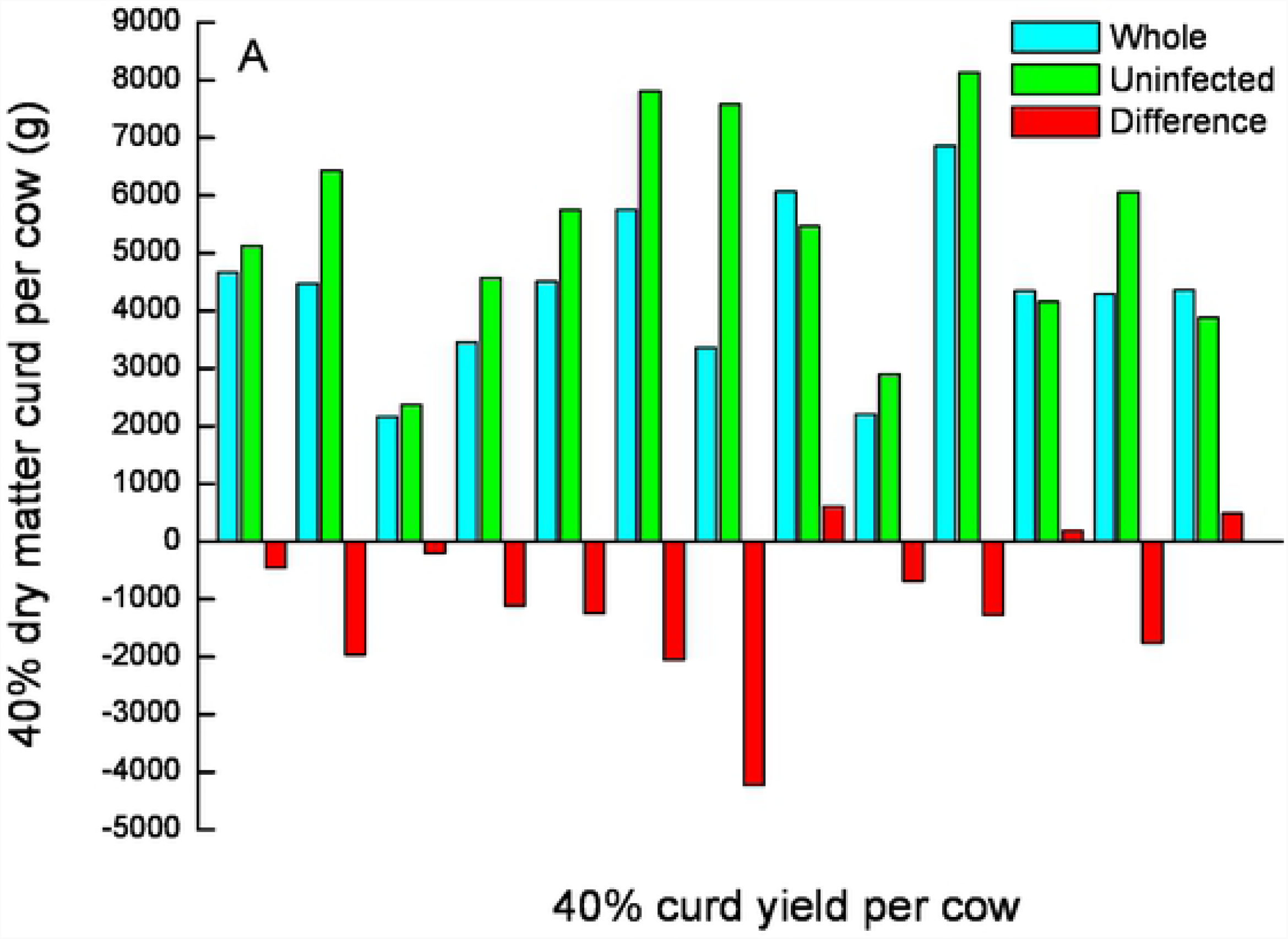
Predicted dry matter of 40% curd of milk from post infected with *E. coli* (PIEc) cows.

**Figure 4b.**
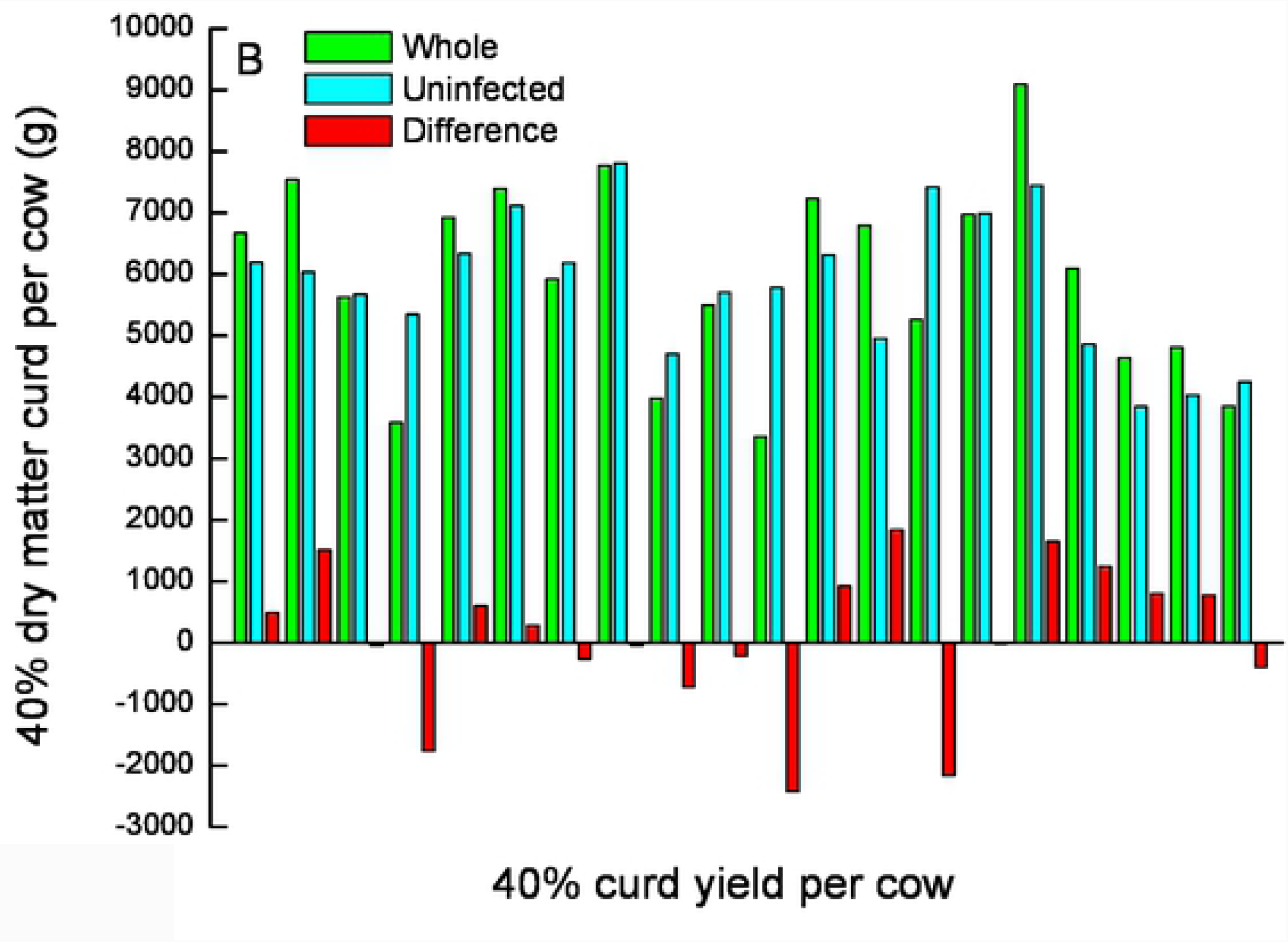
Predicted dry matter of 40% curd of milk from *Streptococcus dysgalactiae* infected cows.

**Figure 4c.**
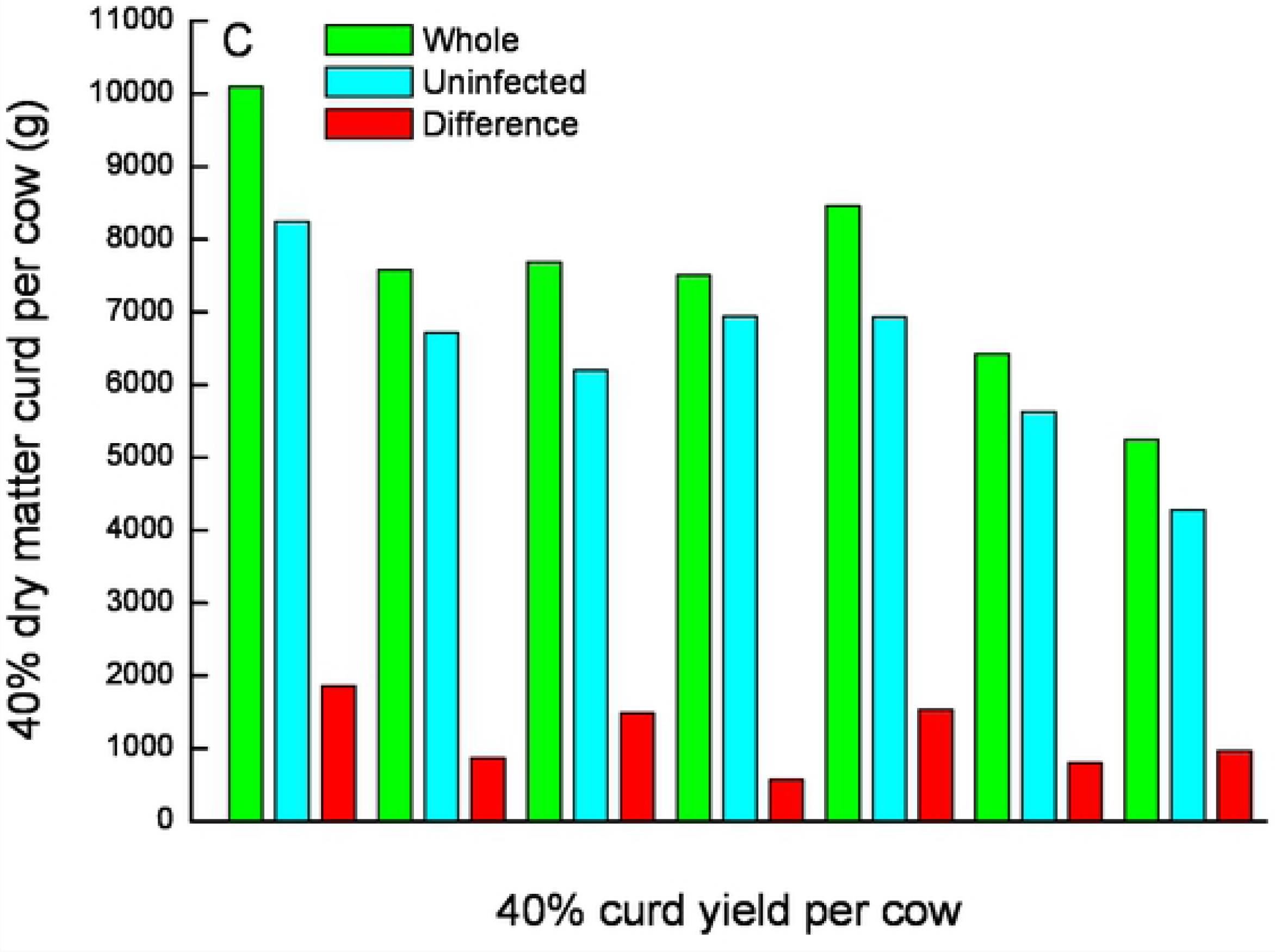
Predicted dry matter of 40% curd of moil from coagulase negative staphylococci cows.

**Figure 5.** Correlation coefficient (R^2^= 0.4) between logSCC in the whole cow milk (healthy and inflamed glands) and curd weight, calculated according to the equation: (8.99 × curd firmness) + 74.2 (see text).

In the milk of all healthy glands log SCC was <5.2 (<2×10^5^ cells/mL) whereas in the milk of all the inflamed glands, regardless of bacteria species, log SCC was > 6.0 (>1×10^6^ cells/mL) (Table 3). The percentage of total leucocytes (CD18^+^) and PMN were significantly higher (*P<0.001*) in all the milk of the inflamed glands (∼85% and ∼70%, respectively) and had a direct influence on the cow’s milk and it was not proportional to milk volume (Table 3). The percentage of macrophages (CD14^+^) was higher (5-9%) in milk of the inflamed glands but significant only in PIEc and *S. dysgalactiae*. CD4^+^ and CD8^+^ T-lymphocytes were ∼4% with no differences between healthy and inflamed glands. Nevertheless, the number of all leucocyte’s types were significantly higher in all inflamed glands owing to higher SCC.

**Table 3.** LogSCC and percent of total leucocytes (CD18), polymorphonuclear (PMN), lymphocytes bearing CD4 or CD8 and microphages (CD14) of uninfected (Healthy), infected and the whole cow’s milk (mixture of healthy glands and the infected one) according to infecting bacteria. Values within rows with no common superscript differ significantly (*P* < 0.05).

|  |  | logSCC | CD18 (%) | PMN (%) | CD4 (%) | CD8 (%) | MØ (CD14) |
| --- | --- | --- | --- | --- | --- | --- | --- |
| PIEc | Healthy | 5.10±0.1<sup>c</sup> | 59.4±6.5<sup>b</sup> | 47.2±5.7<sup>b</sup> | 4.3±1.7 | 3.9±1.2 | 5.4±0.8<sup>b</sup> |
|  | Infected | 6.69±0.1<sup>a</sup> | 88.2±1.3<sup>a</sup> | 69.3±4.1<sup>a</sup> | 3.8±0.8 | 3.4±0.6 | 8.5±1.7<sup>a</sup> |
|  | Whole | 6.21±0.2<sup>b</sup> | 78.9±5.1<sup>a</sup> | 61.3±5.4<sup>a</sup> | 4.5±1.5 | 3.6±1.0 | 5.6±1.0<sup>b</sup> |
| Strep. | Healthy | 5.13±0.1<sup>c</sup> | 62.8±6.6<sup>b</sup> | 53.7±5.2<sup>b</sup> | 3.1±1.1 | 2.5±0.9 | 3.6±0.4 |
|  | Infected | 6.75±0.1<sup>a</sup> | 90.5±1.0<sup>a</sup> | 77.9±3.7<sup>a</sup> | 3.6±1.5 | 3.1±1.2 | 4.9±0.7 |
|  | Whole | 6.23±0.1<sup>b</sup> | 80.6±4.5<sup>a</sup> | 71.3±3.7<sup>a</sup> | 3.0±0.9 | 2.7±0.8 | 3.5±0.4 |
| CNS | Healthy | 4.78±0.1<sup>c</sup> | 55.6±6.0<sup>b</sup> | 46.7±9.0<sup>b</sup> | 3.7±1.8 | 3.6±0.9 | 2.3±0.4<sup>b</sup> |
|  | Infected | 6.29±0.2<sup>a</sup> | 85.4±2.4<sup>a</sup> | 68.7±5.3<sup>a</sup> | 4.9±1.8 | 3.7±1.3 | 5.1±1.8<sup>a</sup> |
|  | Whole | 5.56±0.1<sup>b</sup> | 72.7±6.1<sup>a</sup> | 61.9±6.3<sup>a</sup> | 3.9±1.1 | 4.0±0.9 | 2.0±0.4<sup>b</sup> |

PIEc – post infected by *E. coli*
Strep. – *S. dysgalactiae*
CNS – coagulase negative staphylococci

## Discussion

The current study aimed to measure the influence of the milk of subclinically inflamed mammary glands with different bacteria or post infected glands on the whole cow’s milk quality for the cheese industry and to suggest different parameters for milk payment. The major findings of the present study indicate that inflammation or infection by bacteria such as Streptococci or *E. coli* have a deleterious effect on milk composition and processing related properties, while the effect of bacteria such as CNS is moderate. Drying-off a gland during lactation decreases the overall milk yield of the cow. On the other hand, the milk of affected mammary gland, although representing a minor portion of the whole gland milk yield, may still have a significant negative influence on the quantity and quality of the cheese as was demonstrated by our team [17]. On the gland level, it is acceptable that intramammary infections by different bacteria result in reduction of milk quality and its suitability for maximizing cheese yield. As reported, the damage is tied to the bacteria species with major losses due to the presence of *Staphylococcus aureus* [18], CNS [10,19], *Streptococci* [10,20] and *E. coli* and in particular PIEc [12]. This observation? is intriguing because 1. It depends on the bacteria species, and 2. The effect is observed immediately after mixing of the milk from the affected gland with the healthy ones, i.e. it doesn’t require incubation time, as is the case of milk storage in the bulk tank [4]. The present results clearly demonstrate that not separating the milk from most of the PIEc glands and from ∼ 25% of the glands infected with *S. dysgalactiae* result in lower cheese production.

The findings that different bacteria cause dissimilar damage to the milk showed that clotting parameters are highly affected by hydrolysis of casein[17]. Fleminger et al. [20,21] reported that a low molecular mass fraction isolated from the proteose-peptone preparation of milk, “fraction E”, a casein-derived peptide of 13 kDa rich in phosphorus residues, inhibited milk coagulation. Moreover, fraction E content increased substantially in milk from glands infected with *E. coli* and *S. dysgalactiae*, and during further storage of the milk. It was suggested that chelation of Ca by fraction E was involved in the inhibitory mechanism. This mechanism can explain the immediate negative effect on the coagulation properties of milk from inflamed glands. Although, not tested in the current study, milk is usually stored for 24-48 h before processing. Thus, fraction E-specific enzymes produced by *S. dysgalactiae* that cleave β-casein at a Val-Val amino acid [20] and maybe other enzymes such as plasmin, can have longer negative effects during storage, which increase the justification of discarding this gland’s milk.

Thus, in dairies that produce only cheese and/or fermented products, segregation of the low curd firmness milk from the inflamed glands will end in overall increased product yield as was suggested in several publications on segregating milk for increased cheese yield [15,22]. Moreover, in the eyes of the consumers, guarantee of using only milk that is milked from non-inflamed glands can increase the trust between the dairy and consumers. In order to generate a mechanism where both the farmer and the dairy will benefit from increased product yield, milk of inflamed cow/gland that negatively influence the cow/bulk should be detected. As shown in the present study and others, not every infection and an increase in SCC has or is correlated with negative impacts on product yield [11,23,24]. Because the larger negative impact is related to the cheese industry, a coagulation parameter is probably the most important parameter to be tested before accepting the milk for cheese production. In the Parmigiano Reggiano production routine, coagulation parameters and cow’s udder health, especially in relation to clinical mastitis, are routinely tested and the milk of suspicious cows is not milked into the bulk milk tank [25,26].

In recent years our team has introduced and presented experimental results on the potential on-line separation of milk for cheese manufacturing according to its coagulation properties without any addition or alteration of the milk [15,27-29]. The main results have shown that the level of milk constituents changes throughout the milking session along with the milk’s coagulation properties. Installing the Afilmilk on-line milk channeling system (Afimilk MCS, Afimilk, Afikim, Israel) enables separation of milk according to its measured coagulation properties (Afi-CF) and makes it possible to divert part of the milk into a designated tank and thus to maximize cheese yield [15,29]. Despite the need for higher volumes of water and additional labor to handle, extra cleaning and hauling the milk of the AfiMilk MCS system, economical calculations performed, show a major saving in the total energy consumption owing to the higher cheese yield [30]. The system is operating routinely in a few tens of farms in Israel and Italy, where the dairies process the milk into a variety of hard and soft cheeses.

Currently, in many countries BMTSCC thresholds of 4×10^5^ cells/mL is still under discussion, with the question of how much further pressure should be directed to decrease the BMTSCC is being left open, because there are no clear-cut research results that show what the influence of the further reduction in SCC are. However, no correlation was found between curd firmness and rennet clotting time and cheese yield when BMTSCC ranged from 6.4×10^4^ to 5.97×10^5^ cells/mL, suggesting that up to BMTSCC thresholds of 4×10^5^ cells/mL, the quality of milk does not correlate with SCC since the milk in the tank represents many individual cows [5].

Agriculture in general and dairy farming in particular, as is focused in this study, should guarantee consumers and workers health, as well as animal health and welfare - all under cost-effectiveness. Due to intensive studies over the years, new standards and regulations led to improved products quality by reducing hazard for both humans and livestock and animal welfare. Nevertheless, not every scientific finding and economical investment can or should be applied. In most of the intensive dairy operations, the production is oriented to two major lines: the farms, where the cow is a milk production unit and the dairy, where the bulk milk is the raw material for the production system. Therefore, the decisions and control of each individual cow, if it to be milked into the bulk tank or not regarding its health condition - inflamed and/or infected udder, relies on the bacteria species and its effects. When the mammary infecting bacteria are not particularly pathogenic to humans, or the BTSCC does not reach the rejection level, or and milk price is not penalized, the decision to milk such cow is therefore not trivial. At the same time the dairy industry must recognize that raw milk of higher value needs to be paid extra. Thus, it should price raw milk according to its value for the end product, i.e., as long as the quantity and quality of the end product is not influenced, the price of the milk should remain the same.

## Conclusions

The major findings of the present study indicate that different mastitis causing bacteria species inflict dissimilar damaging effects on the milk. Inflammation or infection by bacteria such as Streptococci or *E. coli* have a deleterious effect on milk composition and processing related properties, while the effect of bacteria such as CNS is moderate. Therefore, the influence of the milk of subclinically inflamed mammary glands with different bacteria species on the whole cow’s milk quality for the cheese industry suggest different parameters for milk payment. As shown in the present study, not every infection and an increase in SCC has or is correlated with negative impacts on product yield. In order to generate a mechanism where both the farmer and the dairy will benefit from increased product yield, milk of inflamed cow/gland that negatively influence the cow’s/bulk tank milk should be detected and a decision should be made whether or not to milk it into the bulk milk tank. The dairy industry must recognize that raw milk of higher value needs to be paid extra. Thus, it should price raw milk according to its value for the end product. Milk that increases product yield should receive a bonus, but if it has no influence on yield, the milk should receive the target price.

